# Clarifying the biological and statistical assumptions of cross-sectional biological age predictors

**DOI:** 10.1101/2023.01.01.522413

**Authors:** Marije H. Sluiskes, Jelle J. Goeman, Marian Beekman, P. Eline Slagboom, Hein Putter, Mar Rodríguez-Girondo

**Affiliations:** Medical Statistics, Department of Biomedical Data Sciences, Leiden University Medical Center, Leiden, the Netherlands; Molecular Epidemiology, Department of Biomedical Data Sciences, Leiden University Medical Center, Leiden, the Netherlands; Max Planck Institute for the Biology of Ageing, Cologne, Germany

**Keywords:** Aging, biological age, aging rate, aging clocks, metabolome

## Abstract

There is variability in the rate of aging among people of the same chronological age. The concept of biological age is postulated to capture this variability, and hence to better represent an individual’s true global physiological state than chronological age.

Biological age predictors are often generated based on cross-sectional data, using biochemical or molecular markers as predictor variables. It is assumed that the difference between chronological and predicted biological age is informative of one’s chronological age-independent rate of aging Δ.

We show that the most popular cross-sectional biological age predictors—based on multiple linear regression, the Klemera-Doubal method or principal component analysis—rely on the same strong underlying assumption, namely that a candidate marker of aging’s association with chronological age is directly informative of its association with the aging rate Δ. We call this the identical-association assumption and prove that it is untestable in a cross-sectional setting. Using synthetic data, we illustrate the consequences if the assumption does not hold: in such scenarios, there is no guarantee that the weights that a cross-sectional method assigns to candidate markers are informative of the underlying truth. Using real data we illustrate that the extent to which the identical-association assumption holds is of direct practical relevance for anyone interested in developing or interpreting cross-sectional biological age predictors.

## Introduction

Individuals of the same chronological age show considerable variation in the rate at which they age: while some enjoy long and healthy lives, others experience early-onset functional decline, suffer from a range of diseases and die young [Partridge et al. 2018]. This variability gave rise to the idea that, in addition to a chronological age, individuals also possess a biological age [Benjamin 1947, Comfort 1969]. This biological age should be an accurate reflection of one’s position on their life-course: when biological age exceeds chronological age this is indicative of accelerated aging (marking a higher physiological vulnerability, lower lifespan expectancy and increased risk to develop (multi)morbidity), the reverse of slow aging.

The question why, how and how fast we age is not only of biological interest, but has direct societal relevance. The enormous increase in average human lifespan that has been observed throughout most of the world in the last centuries has not been matched by an equal increase in healthspan (life years spent in health) [Crimmins 2015, Partridge et al. 2018]. This has led to a global healthcare burden, which is expected to only increase in the decades to come [He et al. 2016]. Measuring biological age could contribute to identifying individuals most at risk and helping them with targeted interventions. In addition, a better insight in the processes that underlie aging might help in designing interventions to slow down, delay or even reverse aging.

Biological age is latent: it cannot be directly measured, which complicates a direct evaluation of predictions. However, there is consensus that biological age contains information on aging above and beyond chronological age [Baker III and Sprott 1988, Jylhävä et al. 2017]. We call this chronological age-independent part of biological age the ‘aging rate’ and denote it by the symbol Δ. Hence, we here mean by aging rate Δ only the biological age acceleration or deceleration (i.e., biological age conditional on chronological age). In line with this consensus, predictions of biological age are generally evaluated by checking if the chronological age-independent part of a prediction, denoted by 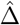, is associated with time-to-death or other outcomes that are known to be measurable physiological outcomes representing the aging process (e.g., grip strength, frailty or cognitive function), in a model adjusted for chronological age.

The aging field is trying to detect (bio)markers indicative of the biological age of individuals, in this paper referred to as ‘candidate markers’ (of biological aging). Such candidate markers of biological aging must be informative of biological age beyond chronological age, i.e., they must be associated with one’s aging rate Δ. Candidate markers can consist of molecular, biochemical, clinical or physiological health data. The earliest attempts to capture biological age made use of a limited number of physiological and biochemical markers [Comfort 1969, Furukawa et al. 1975, Takeda et al. 1982]. More recently, the advent of high throughput bio-molecular technologies has resulted in the development of numerous high-dimensional omics-based age predictors. This renewed interest was initiated by the publication of the Horvath and Hannum DNA methylation (DNAm) age predictors [Horvath 2013, Hannum et al. 2013]. It was soon found that DNAm age predictions are associated with aging above and beyond chronological age [Marioni et al. 2015, Christiansen et al. 2016, Perna et al. 2016]. Since then, various other omics-based age predictors have been developed, e.g. based on IgG glycomics [Kristic et al. 2014], metabolomics [Van Den Akker et al. 2020], proteomics [Tanaka et al. 2018] or transcriptomics [Peters et al. 2015].

Biological age prediction methods, often referred to as ‘aging clocks’, can be divided in two generations. The first-generation prediction methods are based on the association of candidate markers of biological aging with chronological age. These methods hence require cross-sectional data only, where chronological age and candidate markers are measured at a single point in time. The second-generation prediction methods are based on the association of candidate markers with time-to-age-related-event data (as of yet, only time-to-mortality has been considered as outcome of interest). The three most well-known second-generation predictors are PhenoAge [Levine et al. 2018] and GrimAge [Lu et al. 2019], which both use DNAm marker data as (surrogate) predictor variables, and a mortality predictor named MetaboHealth [Deelen et al. 2019], using metabolome data as predictor variables.

Although second-generation epigenetic and metabolomics-based methods outperform first-generation (cross-sectional) methods in terms of their strength of association with time-to-mortality and other aging-related outcomes [Hillary et al. 2020, Maddock et al. 2020, McCrory et al. 2021, Kuiper et al. 2022], cross-sectional methods are still frequently developed, used and debated [Rutledge et al. 2022]. From a practical point of view, the ongoing popularity of cross-sectional methods can easily be explained: cross-sectional data are simply much more abundant than longitudinal (time-to-event) data. Moreover, the predicted aging rates Δ of several recent cross-sectional age predictors were found to be associated with time-to-mortality and the onset of other aging-related outcomes [Marioni et al. 2015, Christiansen et al. 2016, Van Den Akker et al. 2020, Tanaka et al. 2020].

The general consensus in the field therefore seems to be that even though cross-sectional biological age predictors are suboptimal, they still capture some signal related to biological aging, and can therefore still be of value. Nevertheless, how and under which assumptions they can capture this signal is not clear, neither from a statistical nor from a biological point of view. We believe that the statistical assumptions underlying these cross-sectional methods, and the consequences if they are not met, must be known and well understood for aging researchers to evaluate whether it makes sense use to such an approach. Lack of understanding of the assumptions and limitations of any prediction method can hamper progress in the field of biological age prediction and in the identification of relevant markers of aging. Though certain aspects of various cross-sectional methods have been sporadically criticized before (discussed in more detail in the next section), to the best of our knowledge an in-depth discussion of the key assumption that all cross-sectional approaches—often implicitly—rely on does not yet exist.

With this paper we attempt to fill that gap by considering this matter from several angles. We start by providing a comprehensive overview of the most popular cross-sectional biological age prediction methods. We discuss the assumption they all rely on, namely that any marker’s association with chronological age is directly informative of its association with the age-independent part of the difference between predicted and chronological age, denoted by Δ. We call this the identical-association assumption and provide a theoretical result why this assumption is untestable. To illustrate the consequences in settings where this assumption does not (fully) hold, we use two synthetic data examples. Finally, we use real data to illustrate that caution must be taken when using cross-sectional data to predict biological age. With this we hope to increase awareness that all cross-sectional methods that either directly or indirectly rely on candidate markers’ correlation with chronological age may be superfluous, and in any case should not be be used without carefully reflecting beforehand on the assumptions these methods make.

## Methods

### Overview of cross-sectional statistical approaches

By far the most popular statistical approach to estimate biological age (*B*) is to perform multiple linear regression (MLR) on cross-sectional data: chronological age (*C*) is taken as the outcome variable and regressed on a set of candidate markers of biological aging (*X*) that were measured at the same time as chronological age. Then the model’s predicted chronological age is considered to be informative of one’s biological age: 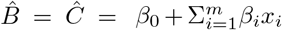, where *m* represents the number of candidate markers included in the regression and *x* represents a single marker. In this method predictions for the aging rate Δ are generally defined as the resulting residuals after regressing predicted biological age (i.e., 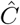) on chronological age. Hence, the residuals of the chronological age model are considered to be informative of Δ. This approach is used with both low- and high-dimensional markers.

The MLR approach does not follow from an underlying model of biological age. It fully relies on a model that predicts chronological age to be indicative of the aging rate Δ. For this to work it must hold that markers that are correlated with chronological age are also correlated with Δ, and vice versa. In fact, it is implicitly assumed that the higher the correlation with chronological age (in a multivariable model, so adjusting for all other included markers), the stronger it is correlated with Δ. Markers that are insignificant predictors of chronological age are assumed to be insignificant predictors of Δ.

Although the MLR approach is the most often-used cross-sectional approach, it has been criticized for various reasons. It suffers from inherent methodological problems, such as regression to the mean (fitted values regress towards the sample’s mean age such that biological ages calculated for those younger than the sample mean age tend to be too high and for those older, too low) and the so-called ‘biomarker paradox’ (a (bio)marker that perfectly correlates with chronological age is useless in estimating biological age) [Ingram 1988, Hochschild 1989]. The biomarker paradox is more than a mere theoretical danger: with epigenetic biological age predictors, in principle a near-perfect chronological age predictor can be developed, as long as the sample size is large enough [Zhang et al. 2019]. In such a case all signal related to biological aging would be lost. This paradox therefore illustrates the peculiarities that arise when the residuals of a linear regression are interpreted as meaningful quantities in their own right, while in the model formulation those residuals are per definition nothing but noise.

Alternative cross-sectional approaches have been proposed in an attempt to overcome some of these methodological issues. The most notable alternatives are principal component (PC)-based methods and the Klemera-Doubal (KD) method [Klemera and Doubal 2006]. PC-based methods transform candidate markers to a set of uncorrelated principal components [Nakamura et al. 1988, Jee and Park 2017, Jia et al. 2017, Pyrkov et al. 2018]. Most of the times, first a pre-selection of candidate markers is made based on how strongly each individual marker is correlated with chronological age. Often, the first principal component of this subset of variables is found to be correlated with chronological age and is hence interpreted as an ‘unscaled’ or ‘standardized’ biological age score *BS*. This score is sometimes transformed to an age-scale based on the mean and standard deviation of chronological age (*μ_C_* and *σ_C_*) in the training sample: *B* = *BS* * *σ_C_* + *μ_C_*.

The Klemera-Doubal method [Klemera and Doubal 2006] uses a reversed regression approach (regressing each candidate marker on chronological age). In contrast to the above methods, the KD method is based on an explicit underlying model of biological age. It assumes that the relation between biological age and chronological age can be expressed by *B* = *C*+Δ. Each marker *x* is governed by *B* but is also affected by random fluctuations. Assuming a linear relation between marker *x* and biological age, *x* equals *β*_0_ + *β*_1_ * *B* + *ϵ*. This can also be expressed as *x* = *β*_0_ + *β*_1_ * (*C* + Δ) + *ϵ*. That the coefficient *β*_1_ is the same for *C* and Δ is a key assumption of the Klemera-Doubal method: in their model, a marker’s strength of association with chronological age is directly informative of its association with Δ. A biological age prediction is obtained by taking a linear combination of all included markers, each of them weighted in terms of the estimated slopes and residual variances resulting from the reversed regressions.

Though in certain settings the Klemera-Doubal method has been found to outperform MLR- and PC-based methods [Levine 2013], extending the method to high-dimensional settings is not straightforward, since it assumes that all included markers are functionally uncorrelated. Therefore the KD method is primarily used in low-dimensional settings [Cho et al. 2010, Jee and Park 2017, Mitnitski et al. 2017], or prior to applying the KD method principal component analysis is used to obtain a set of lower-dimensional markers [Levine 2013, Earls et al. 2019]. The limitations of the alternative cross-sectional approaches might explain the continued popularity of the MLR approach in high-dimensional settings. In a recent review of omics-based biological age predictors the Klemera-Doubal method is not mentioned and PC-based methods play a minor role [Rutledge et al. 2022].

### Reflection on the assumption underpinning cross-sectional biological age predictors

The cross-sectional methods described above share a common assumption, namely that a candidate marker’s strength of association with chronological age is identical to its strength of association with one’s aging rate (the difference between biological and chronological age) Δ. So by using one of the above cross-sectional methods for biological age prediction it is assumed that the traits most strongly associated with chronological age are the ones most informative of Δ. If a marker changes with chronological age irrespective of relevant changes in Δ, or vice versa, the assumption is not met.

For ease of reference, we henceforth refer to this assumption as the *identical-association assumption*. The KD method explicitly makes this assumption. The MLR approach implicitly relies on it (here it concerns ‘adjusted’ association in a multivariable setting). Markers with high absolute coefficient values will have a strong effect on the resulting chronological age prediction 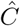, which is considered equal to biological age prediction 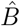. For the PC-based approaches this assumption is used when making a pre-selection of markers prior to finding the principal components, since only variables significantly correlated with chronological age are selected. It is therefore not surprising that the first principal component is often found to be correlated with chronological age: the variables were selected to share this common source of variance.

There are different degrees to which the identical-association assumption might hold in real data. For any set of candidate markers of biological aging, one can roughly distinguish four possible scenarios. The first scenario is that the identical-association assumption holds. If one would then plot the true association of markers with chronological age against their true association with aging rate Δ, one would end up with a plot as given in the top left panel (A) of Figure 1. (There are of course many ways to define ‘association’ – since we do not want to assume a specific model, we deliberately keep this term vague. The plots are therefore conceptual representations of the four scenarios.) As mentioned, the Klemera-Doubal method explicitly makes this assumption, as it assumes an identical regression coefficient (effect size) for chronological age and aging rate Δ and no other sources of shared variance. In this first scenario it would make perfect sense to use a cross-sectional prediction method. The second scenario (shown in panel B of Figure 1) is one in which the opposite of the identical-association assumption holds: the stronger a marker is positively associated with chronological age, the stronger it is negatively associated with aging rate Δ. This is an unlikely possibility, which is only included such that the four scenarios discussed here are collectively exhaustive. The third scenario (shown in panel C of Figure 1) is that the markers’ strength of association with chronological age is not informative of their association with aging rate Δ at all. In such a scenario, using a cross-sectional prediction method would be useless: the weights that cross-sectional methods give to markers will be based on their association (both strength and direction) with chronological age, but these weights will be completely uninformative of the markers’ association with Δ. The fourth and final possibility (shown in panel D of Figure 1) is that the markers’ strength of association with chronological age is somewhat, but not exactly, informative of their association with aging rate Δ. Of the four scenarios this appears to be the most realistic one.

**Figure 1:**
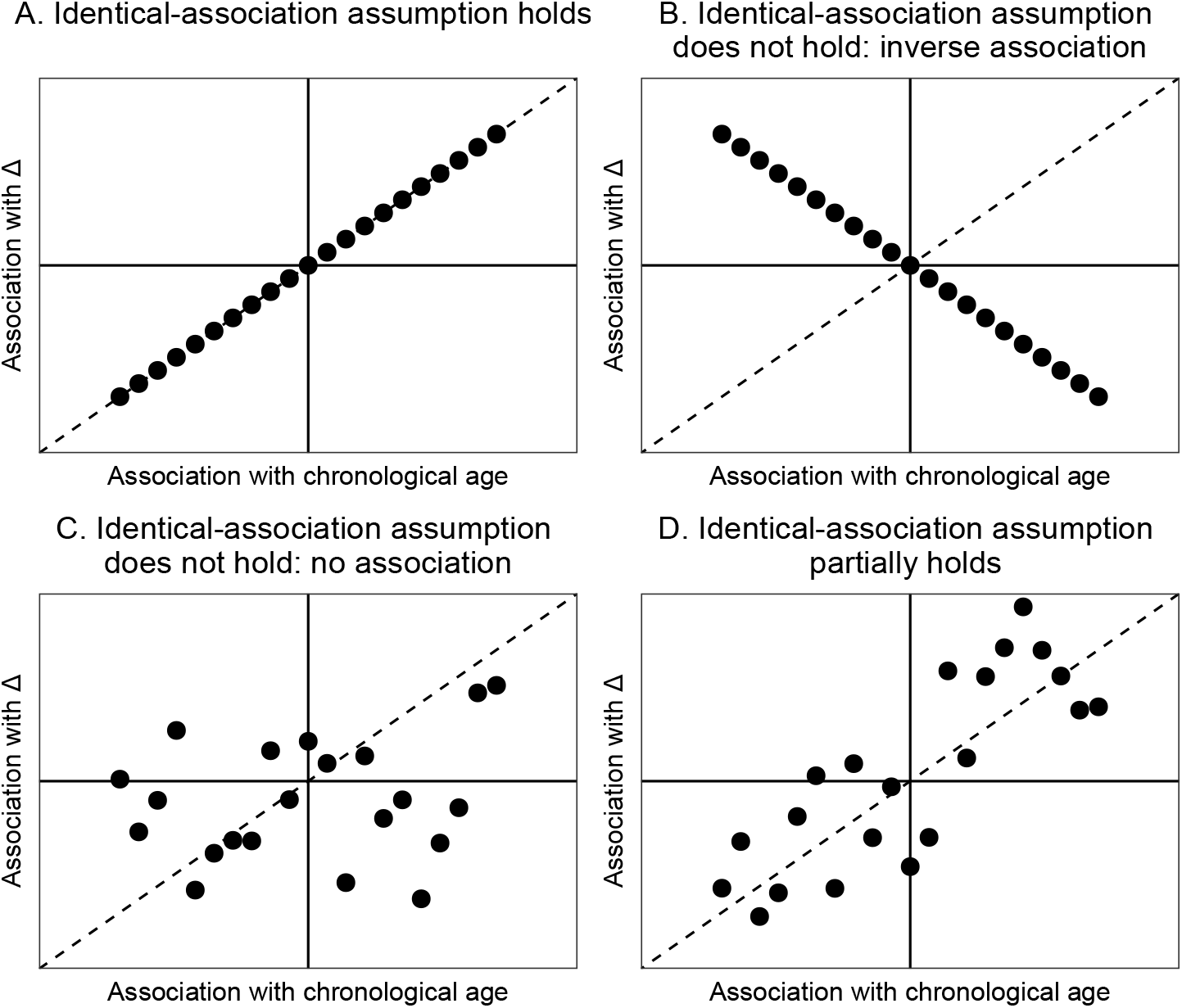
Conceptual visualization of a scenario in which the identical-association assumption holds (A), a scenario in which the inverse relation holds (B), a scenario in which there is no association (C) and a scenario in which the identical-association assumption partially holds (D).

For this fourth scenario it is important to remember that many of the high-dimensional cross-sectional biological age predictors perform some kind of marker selection, either before including them in the model or during the model fitting itself. If one would then only include the variables most strongly correlated with chronological age (i.e., only the edges of Figure 1D would be included, as illustrated in Figure 2), there no longer is a relation between strength of association with chronological age and with aging rate Δ. However, in Figure 2 there still is a relation between the *direction* of the association of the selected markers with chronological age and with Δ. This suggests that in a scenario where the fourth scenario holds and candidate markers of biological aging have been pre-selected, the size of a candidate marker’s association with chronological age will not be informative of its association with Δ, but the sign (positive/negative) of this association will be.

**Figure 2:**
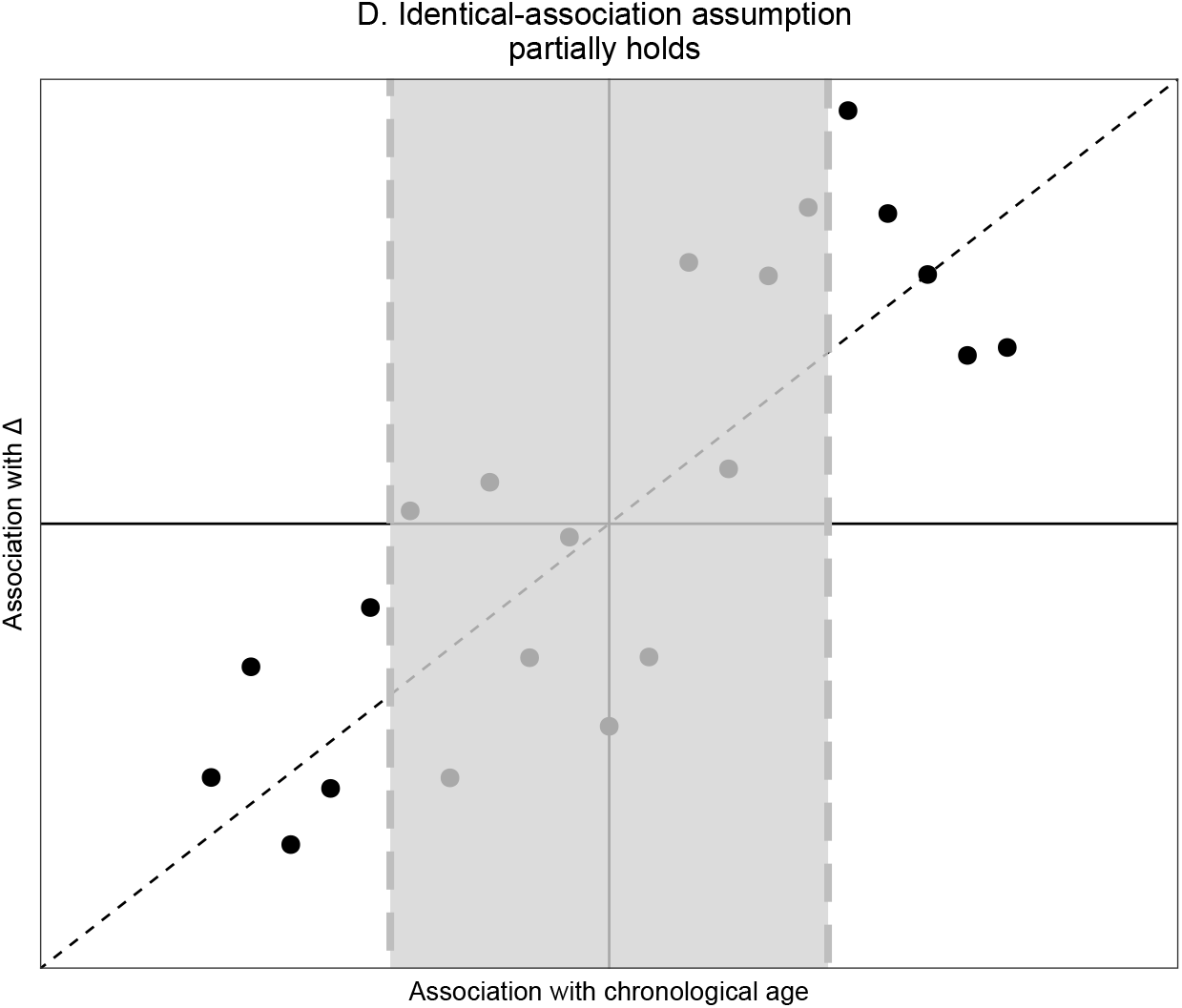
Zoomed-in version of the bottom right panel of Figure 1. If markers are (pre-)selected based on their strength of correlation with chronological age, those in the grey area (i.e., those most weakly associated with chronological age) are not selected.

Which scenario holds in a given data set determines whether or not it makes sense to use a cross-sectional method to predict biological age. Unfortunately, in cross-sectional data the identical-association assumption cannot be proven or disproven, because it is untestable: it is impossible to tell to what extent a marker is associated with aging rate Δ based on its association with chronological age alone. For a formal theorem and proof of the untestability of the identical-association assumption we refer to the Supplementary Materials (Appendix A). An intuitive visualization of the proof is given in Figure 3. It shows correlation Venn diagrams [Ip 2001] for two candidate markers of biological age, *X* and *X*′. The two candidate markers have the same association with chronological age *C*, but where marker *X* shares association with biological age *B*, candidate marker *X*′ has no such association. Since *B* is unobserved, we only have information on the joint distribution of *X* and C, or *X*′ and C, respectively. With respect to this observable variation, the diagrams for *X* and *X*′ are identical. It follows that we cannot distinguish between the true marker *X* of biological age and the false marker *X*′. Hence, if the identical-association assumption does not hold, it is impossible to distinguish true markers of Δ from false ones. Using cross-sectional biological age prediction methods, thereby (implicitly) believing in the identical-association assumption, is therefore based on biological hope or knowledge alone, not on a statistical property of the cross-sectional methods.

**Figure 3:**
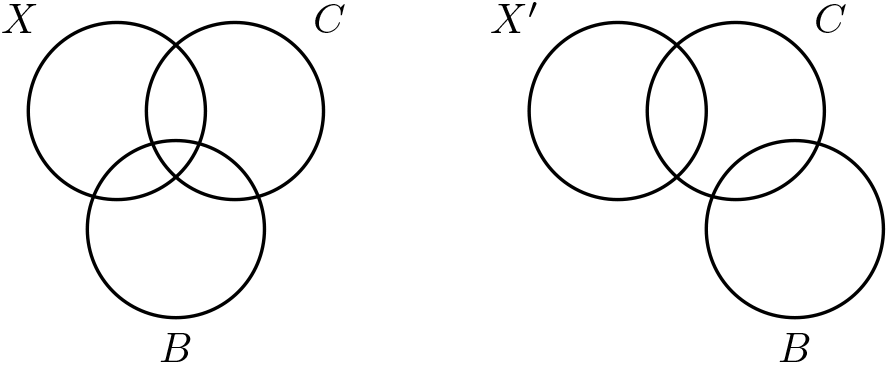
Venn diagrams illustrating the variance shared between biological age (*B*), chronological age (*C*) and the candidate markers of biological aging *X* (true, left diagram) and *X*′ (false, right diagram). Black indicates observed variance; grey unobserved.

## Results

### Two illustrative examples

This section contains two synthetic data examples that illustrate two aspects of the identical-association assumption.

#### Example 1: untestability of the identical-association assumption

We created a synthetic data set with four variables: chronological age *C*, biological age *B*, true marker of biological age *X* and false marker of biological age *X*′. *X* and *X*′ follow the same distribution and have the same strength of correlation with *C*. We based our data generation approach on the type of additive model proposed by Klemera and Doubal [Klemera and Doubal 2006]. We generated *n* observations as follows:

1. Independently generate the following elements:
  - 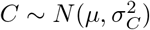;
  - 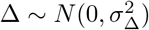;
  - 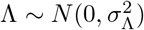;
  - 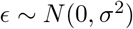;
  - 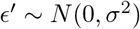.
2. From these elements, construct:
  - *B* = *C* + Δ;
  - *X* = *α* + *β* × (*C* + Δ)+ *ϵ*;
  - *X*′ = *α* + *β* × (*C* + Λ) + *ϵ*′.

We used the following parameter values: *n* = 1000, *μ* = 50, 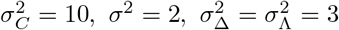, *α* =1, *β* =1.

*X* and *X*′ have the same distribution and the same relation with chronological age, as seen in Figure 4. However, *X* correlates with the individual aging rate Δ while *X*′ does not, as seen in Figure 5. This implies that *X* has useful information on biological age that is not already in chronological age while *X*′ does not. However, in real cross-sectional data Δ is not observed: with respect to their association with the observable variable chronological age these two candidate markers are identical, as can be seen in Figure 4.

**Figure 4:**
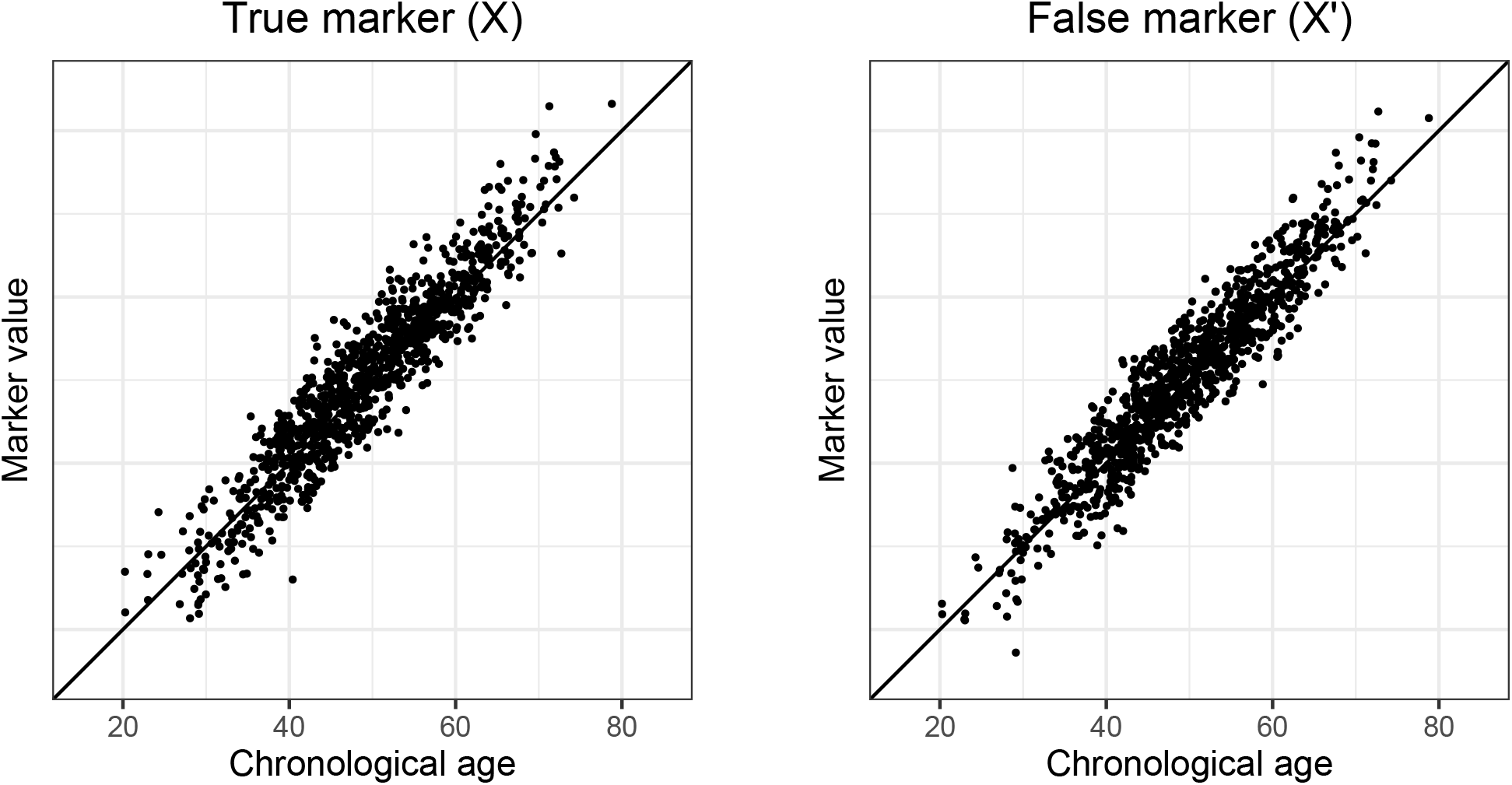
Chronological age plotted against the marker value for true marker *X* and false marker *X*′.

**Figure 5:**
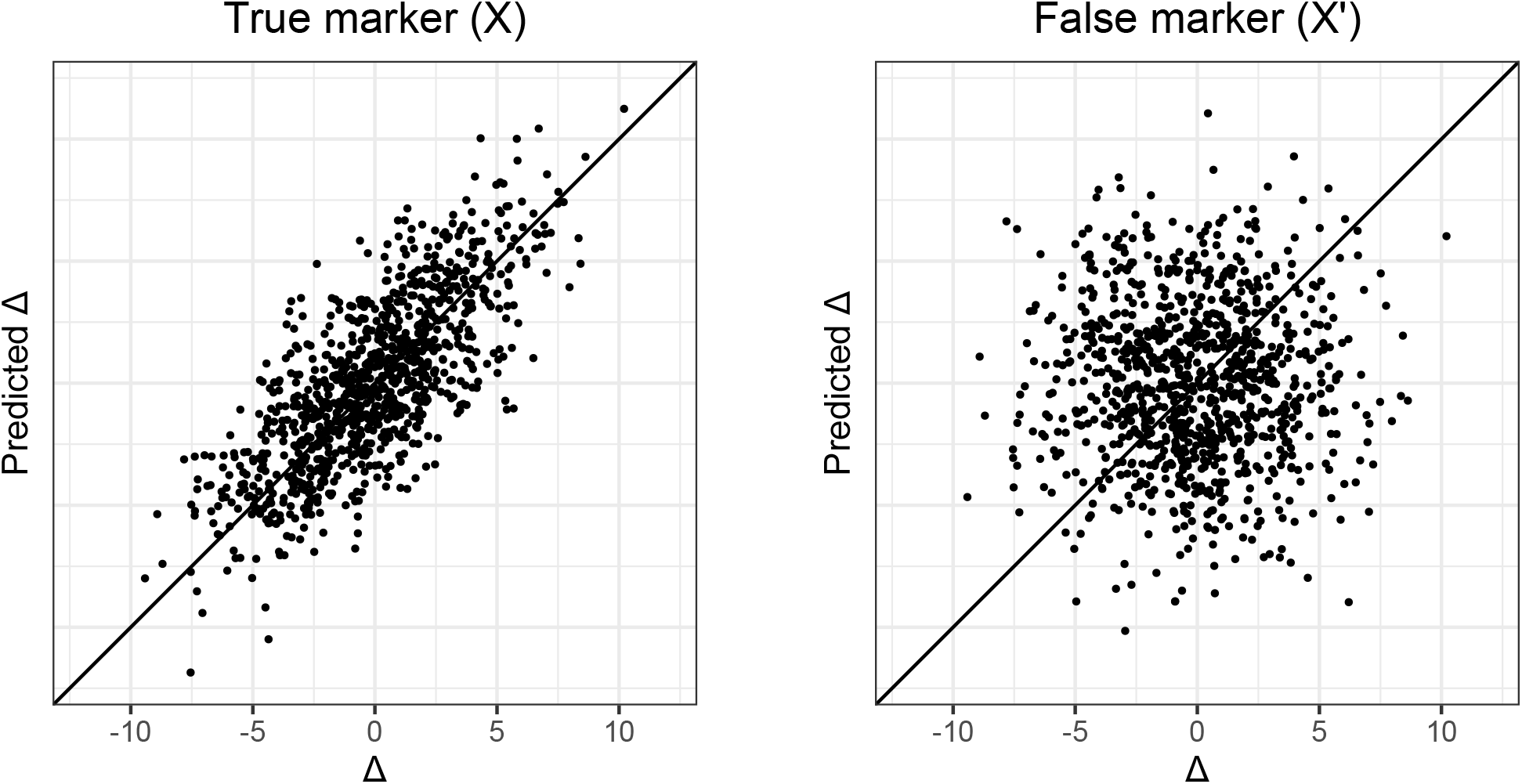
Rate of aging Δ (the difference between true biological and chronological age) plotted against predicted Δ (the resulting residuals after regressing predicted age on chronological age) for true marker *X* and false marker *X*′. The biological age predictions were obtained using linear regression.

Since the observable data (*X, C*) and (*X*′, *C*) are indistinguishable from each other, any method we would apply on either (*X, C*) or (*X′, C*) would assign the same weight to either *X* or *X*′. This holds for the linear regression method, as is clear from Figure 5. It also holds for the Klemera-Doubal method, since that method would assign the same weights to both *X*′ and *X*. Principal components-based methods would not be able to distinguish an informative source of variance (i.e., Δ) from an uninformative source of variance (here denoted by Λ). In fact, no cross-sectional method can distinguish between *X* and *X*′ based on their association with chronological age *C*, because the identical-association assumption is untestable. Therefore, no cross-sectional method can provide evidence that a candidate marker is a truly informative *X* rather than a completely uninformative *X*′.

#### Example 2: consequences of believing in the identical-association assumption under the four different scenarios

The first example illustrated that cross-sectional methods cannot be relied upon to select true markers of the rate of aging Δ. Nevertheless, predicted Δ-values of several cross-sectional age predictors have been found to be associated with time-to-mortality and several other age-related outcomes [Rutledge et al. 2022], albeit often weakly. This can only be the case if a marker’s strength of correlation with chronological age is at least somewhat indicative of its strength of association with true aging rate Δ.

To illustrate this, we generated a possible realization of each of the four conceptual scenarios depicted in Figure 1. We obtained predictions for aging rate Δ using multiple linear regression (MLR) and the KD-method. We know that if the identical-association assumption does not hold, the weights found by MLR and the KD-method are uninformative of a marker’s strength of association with Δ. If the identical-association assumption partially holds, the size of the weights that cross-sectional methods assign to markers will not informative but the signs (positive/negative direction) of these weights still are (Figure 2). We illustrate this by also including a third, ‘naive’ prediction method in this second example, where similar to the MLR approach we took a linear combination of markers. In this third prediction method each marker was assigned the same weight, namely the mean of the MLR coefficients. The sign of each coefficient was kept unchanged, because we generally expected the sign to be correct. We included this third approach to illustrate that if the identical-association assumption does not hold, weights obtained using the MLR or Klemera-Doubal method might result in less accurate predictions than naively assigning each marker the same weight.

For this second example we generated four data sets, *DF_A_, DF_B_*, *DF_C_* and *DF_D_*, corresponding to the scenarios in Figure 1. To keep it simple, each data set has only three markers(*X*_1_, *X*_2_ and *X*_3_), which are associated with chronological age *C* and with aging rate Δ to varying degrees in each of the four scenarios.

We generated *n* observations as follows:

1. Independently generate:
  - 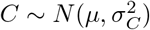;
  - 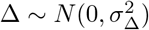;
  - 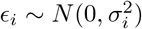.
2. Construct biological age:
  - *B* = *C* + Δ.
3. Construct markers:
  - *X*_1_ = *β*_*C*,1_ × *C* + *β*_Δ, 1_ × Δ + *ϵ*_1_;
  - *X*_2_ = *β*_*C*,2_ × *C* + *β*_Δ, 2_ × Δ + *ϵ*_2_;
  - *X*_3_ = *β*_*C*,3_ × *C* + *β*_Δ, 3_ × Δ + *ϵ*_3_.

Per scenario, the values chosen for *β*_*C*,1_ and *β*_Δ,1_ can be found in Table 1. The following parameter values were used in all four scenarios: *n* = 1000, 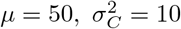 and 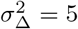. The standard deviation of the errors *ϵ* were chosen such that the relation between the (scaled and centered) three markers and chronological age is the same in in all four data sets (Supplementary Materials, Appendix B). Hence, based on the observable variables alone (*X*_1_, *X*_2_, *X*_3_ and *C*) the four data sets are indistinguishable.

**Table 1:** The coefficients used to construct the markers *X*_1_, *X*_2_ and *X*_3_ for the four different scenarios.

| | $X_1$ | | $X_2$ | | $X_3$ | |
| --- | --- | --- | --- | --- | --- | --- |
| | $\beta_C$ | $\beta_\Delta$ | $\beta_C$ | $\beta_\Delta$ | $\beta_C$ | $\beta_\Delta$ |
| Scenario A | 10 | 10 | 3 | 3 | 5 | 5 |
| Scenario B | 10 | -10 | 3 | -3 | 5 | -5 |
| Scenario C | 10 | 0 | 3 | 10 | 5 | -1 |
| Scenario D | 10 | 9 | 3 | 5 | 5 | 5 |

If the identical-association assumption holds (scenario A), the MLR approach and the Klemera-Doubal approach outperform the equal weights approach (Figure 6: the closer the points are to the diagonal line Δ = predicted Δ, the better the performance of the method). In this case a marker’s association with chronological age is directly informative of its association with rate of aging Δ, so any method that weighs markers according to their strength of correlation with chronological age will do well. In the unrealistic case that a marker’s association with chronological age is inversely related to its association with aging rate Δ (scenario B), all methods will perform badly, as is to be expected (Figure 7). If there is no relation between a marker’s association with chronological age and its association with aging rate Δ (scenario C), the equal weights approach outperforms the two cross-sectional approaches, which appear to capture only noise (Figure 8). In our realization of scenario D, it can be seen that all methods capture some signal (Figure 9). The Klemera-Doubal method does best, but that might not be surprising given the data generation approach, which was based on the type of additive model Klemera and Doubal assume. Interesting is that the same weights approach outperforms the MLR-approach. Naturally, this is just one possible realization: with different values for *β_C_* and *β*_Δ_ the coin could flip in favor of the MLR-method over the equal weights method (and with a different data generation mechanism, possibly also over the KD-method).

**Figure 6:**
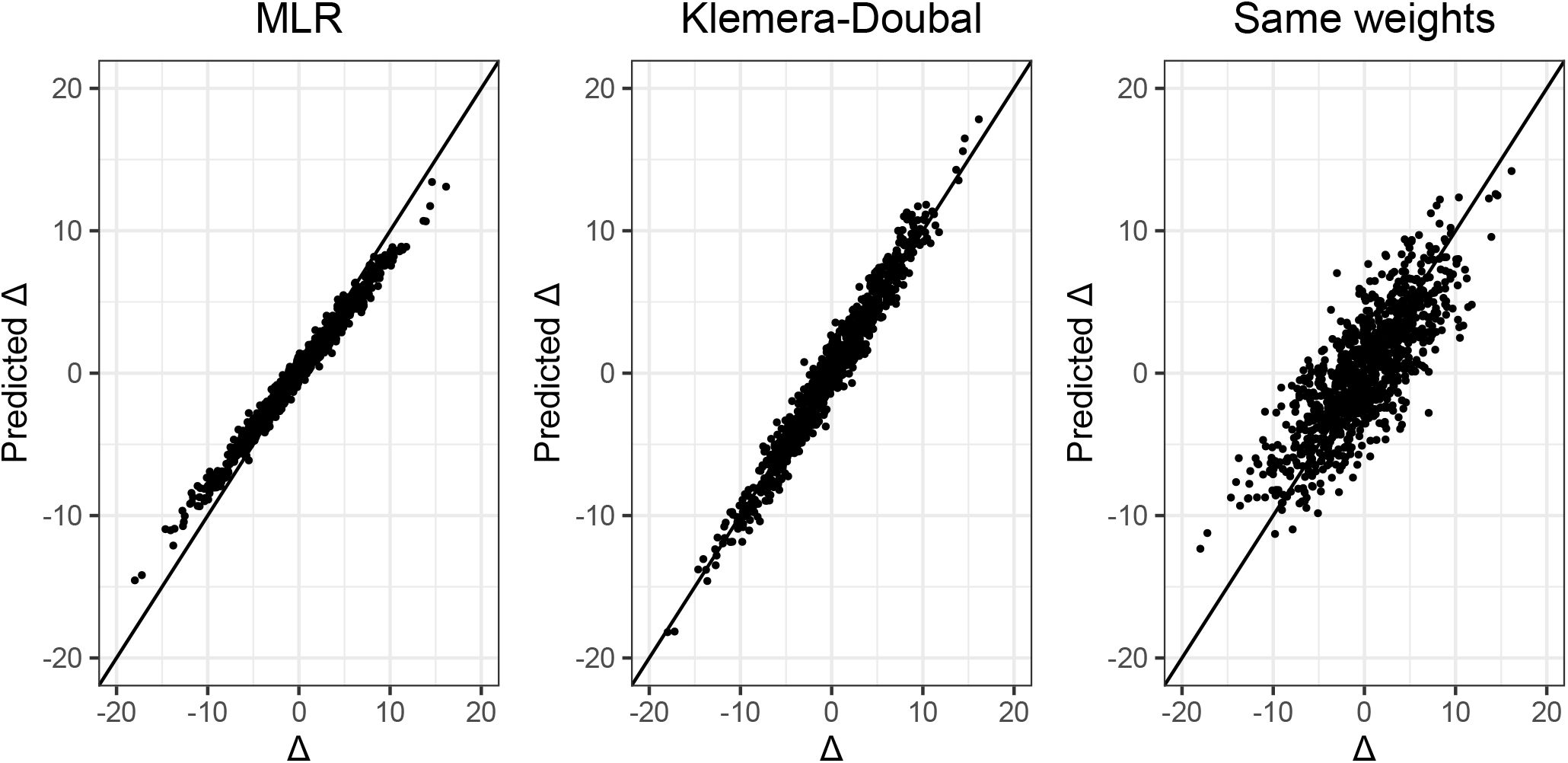
Rate of aging Δ plotted against predicted Δ for the MLR method, the Klemera-Doubal method and the MLR method where each marker is assigned the same weight. Here the identical-association assumption holds (scenario A).

**Figure 7:**
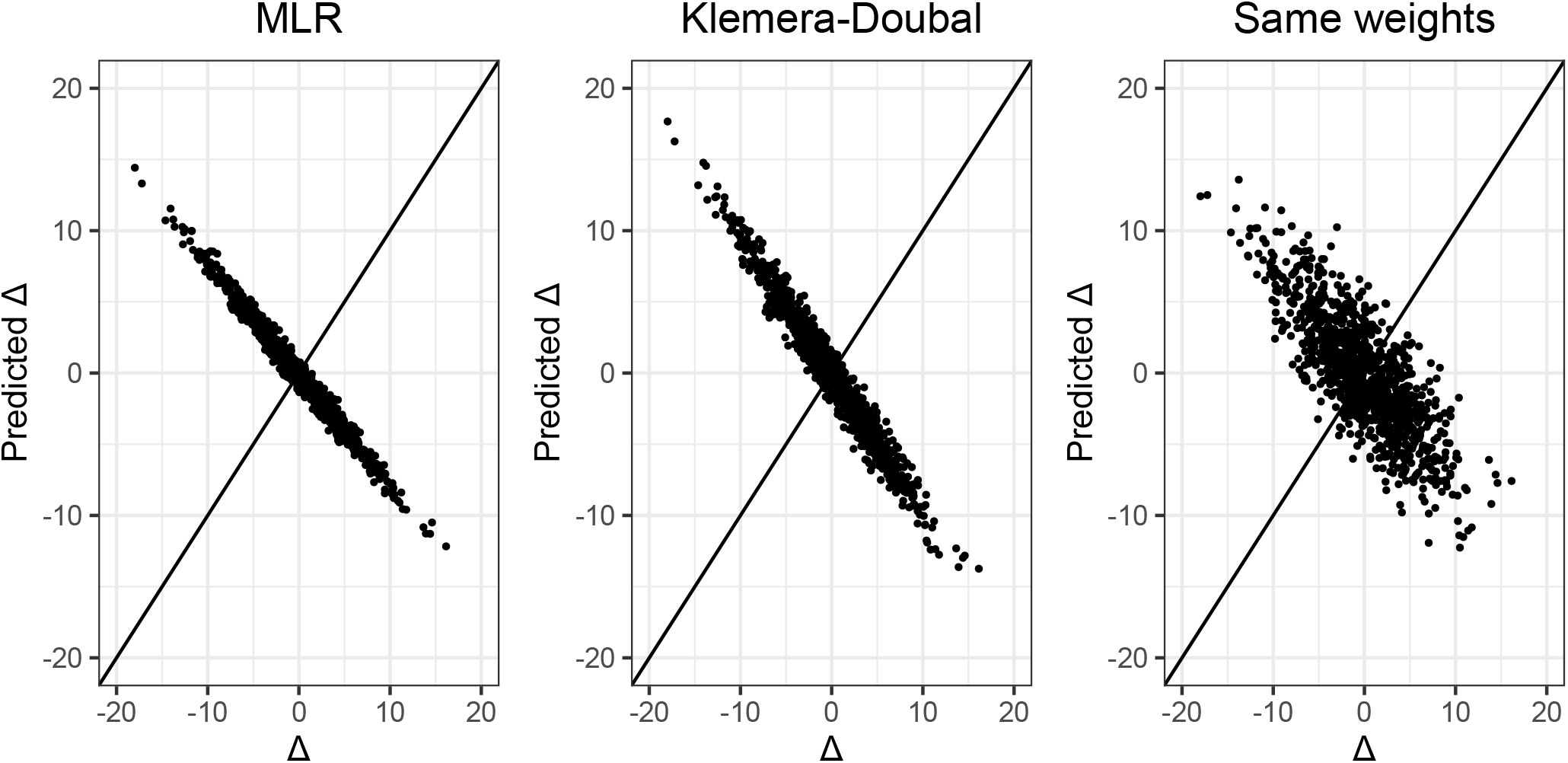
Rate of aging Δ plotted against predicted Δ for the MLR method, the Klemera-Doubal method and the MLR method where each marker is assigned the same weight. Here the identical-association assumption does not hold, but an inverse relation exists (scenario B).

**Figure 8:**
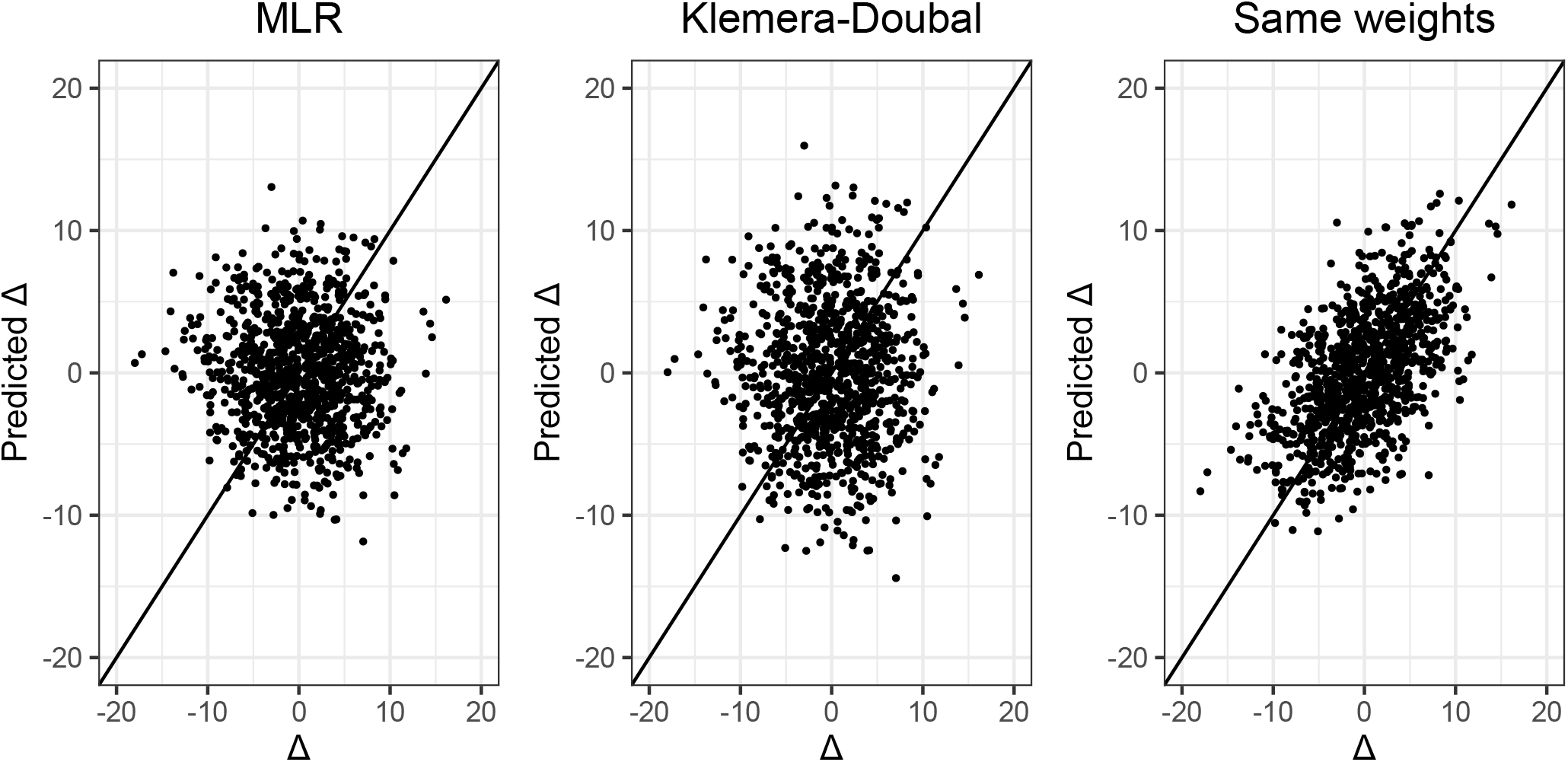
Rate of aging Δ plotted against predicted Δ for the MLR method, the Klemera-Doubal method and the MLR method where each marker is assigned the same weight. Here the identical-association assumption does not hold: there is no association (scenario C).

**Figure 9:**
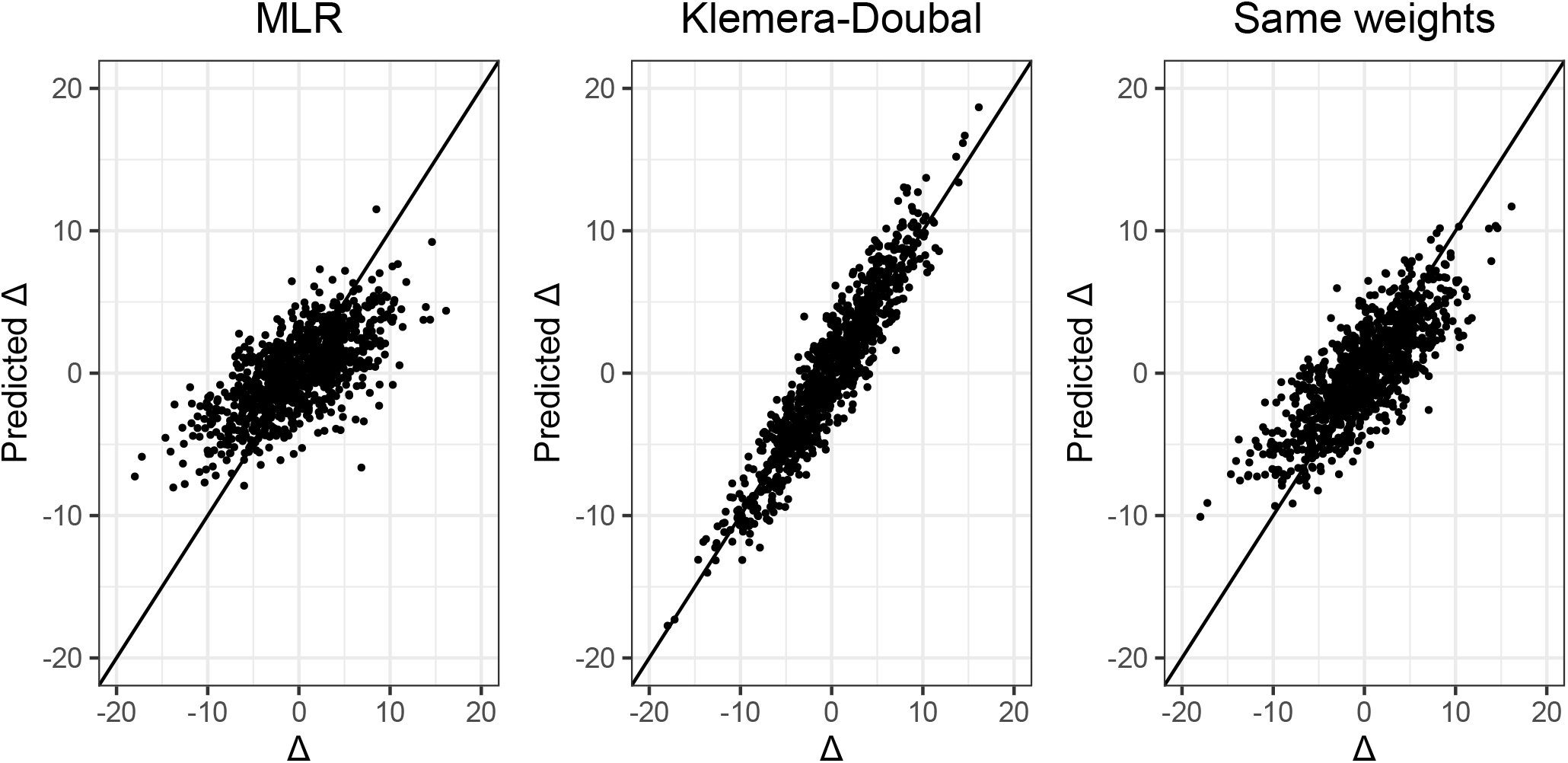
Rate of aging Δ plotted against predicted Δ for the MLR method, the Klemera-Doubal method and the MLR method where each marker is assigned the same weight. Here the identical-association assumption partially holds (scenario D).

### Real data illustration

The insights gained from the synthetic data scenarios are of immediate practical relevance. We illustrate this with a real data illustration.

We used data from the the Leiden Longevity Study (LLS) [Westendorp et al. 2009]. The LLS follows long-lived siblings of Caucasian descent, their offspring and the partners of their offspring. We used data on the offspring and partners (N = 2312). Participants who were lost to follow-up (N = 10) or who had at least one missing metabolite value (N = 37) were excluded. In total 1593 offspring and 674 partners were included, of which 998 men and 1269 women (mean age at inclusion 59.15 years, sd 6.72). Participants were included between March 2002 and May 2006. Registry-based follow-up until November 2021 was available. Median follow-up time was 16.26 years (IQR: 15.31–17.08). 309 deaths were observed. The Medical Ethics Committee of the Leiden University Medical Center approved the study and informed consent was obtained from all participants.

As candidate markers of biological aging we used blood-based metabolic variables. The metabolic variables were quantified using a well-standardized high-throughput nuclear magnetic resonance (^1^H-NMR) metabolomics platform [Soininen et al. 2015, Würtz et al. 2017] of Nightingale Health Ltd. (Helsinki, Finland). Of the more than 200 metabolic variables available, a subset of 59 was selected, previously found to be most reliable and independent [Deelen et al. 2019] and used in various subsequent publications [Van Den Akker et al. 2020, Bizzarri et al. 2022]. Prior to analysis, a small constant was added to all metabolic variables after which they were log-transformed and scaled.

The complete two-generation Leiden Longevity Study has previously been used in two major analyses by our group, constructing biological age predictors (on cross-sectional as well as time-to-event basis) based on the same metabolic variables in much larger data sets. From these studies we observed that the constructed predictors as well as many of the 59 metabolic variables separately were predictive of prospective mortality [Deelen et al. 2019, Van Den Akker et al. 2020].

To illustrate the problems that can arise when using cross-sectional methods to predict biological age, we took a similar approach as in synthetic data example 2: we contrasted an often-used cross-sectional approach to obtain predictions for rate of aging Δ—in this case penalized regression, hereafter denoted by method 1 —with naive methods to obtain predictions for Δ—in this case first selecting metabolites univariately associated with chronological age and then using (unpenalized) multiple linear regression (method 2), a linear combination with either equal weights (method 3) or randomly drawn weights (method 4).

For each of the four methods, predictions for aging rate Δ were obtained as follows. For method 1 we first obtained an age prediction using penalized MLR with a ridge penalty. Using 10-fold cross-validation, the penalization parameter λ was chosen such that the mean cross-validated error was minimized. Chronological age was taken as the outcome variable and all 59 metabolic variables were included as predictor variables. In method 2 we performed (unpenalized) multiple linear regression on a subset of variables correlated with chronological age. 26 of the 59 metabolic variables were significantly correlated with chronological age, using a Bonferroni-corrected significance threshold of 0.05/59 = 8.47 × 10^−4^. For method 3 we again took a linear combination of the 26 metabolic variables significantly correlated with chronological age. Here we assigned each variable same weight, namely the mean of the absolute value of the MLR-coefficients from method 2 (excluding the intercept). Although the coefficients were averaged, the *sign* of each variable’s coefficient was kept, for the same reason as illustrated by Figure 2: it is unlikely that a variable is positively correlated with chronological age but negatively with Δ. Method 4 is a variation on method 3: 1,000 different linear combinations of the same 26 variables were taken, where each variable was assigned a coefficient randomly drawn from a uniform distribution. Similar to method 3, the weights were drawn at random but the signs were kept. For each of the four methods, predictions for Δ were obtained by regressing the the linear combination of metabolic variables (the fitted values) on chronological age and obtaining the residuals.

We then compared the performance of the four methods by scaling the predictions for aging rate Δ obtained using each of the four methods and including them in a Cox proportional hazards (PH) model with time-to-mortality as outcome. This is a common approach to check the validity of Δ-predictions if data on time-to-death is available [Marioni et al. 2015, Zheng et al. 2016, Christiansen et al. 2016, Van Den Akker et al. 2020, Tanaka et al. 2020, Hillary et al. 2020, McCrory et al. 2021]. We used chronological age as the timescale of the Cox PH model and adjusted for sex. Since all predicted Δ-values were scaled prior to inclusion, the higher the coefficient for Δ, the stronger the association with time-to-mortality.

The Cox PH coefficients of the different Δ-predictions (i.e., the effect sizes of the association with prospective mortality) obtained with these four methods are compared in Figure 10. It can be seen that the coefficient for aging rate Δ obtained with method 1 is lower than those of methods 2 and 3: hence, association with time-to-death is weaker. The blue and green lines of methods 2 and 3 are very close to each other: using multiple linear regression (method 2) works just as well as assigning each marker the same coefficient (method 3). The histogram represents the distribution of the 1,000 coefficients obtained by assigning each metabolic variable a randomly drawn weight (method 4), repeated 1,000 times. More than half of the histogram area is to the right of the yellow line of the ridge-based coefficient (method 1), and a substantial part is even to the right of the blue and green lines of methods 2 and 3. These results imply that in this particular setting, the naive methods capture more signal related to prospective mortality than the ‘proper’ cross-sectional method 1.

**Figure 10:**
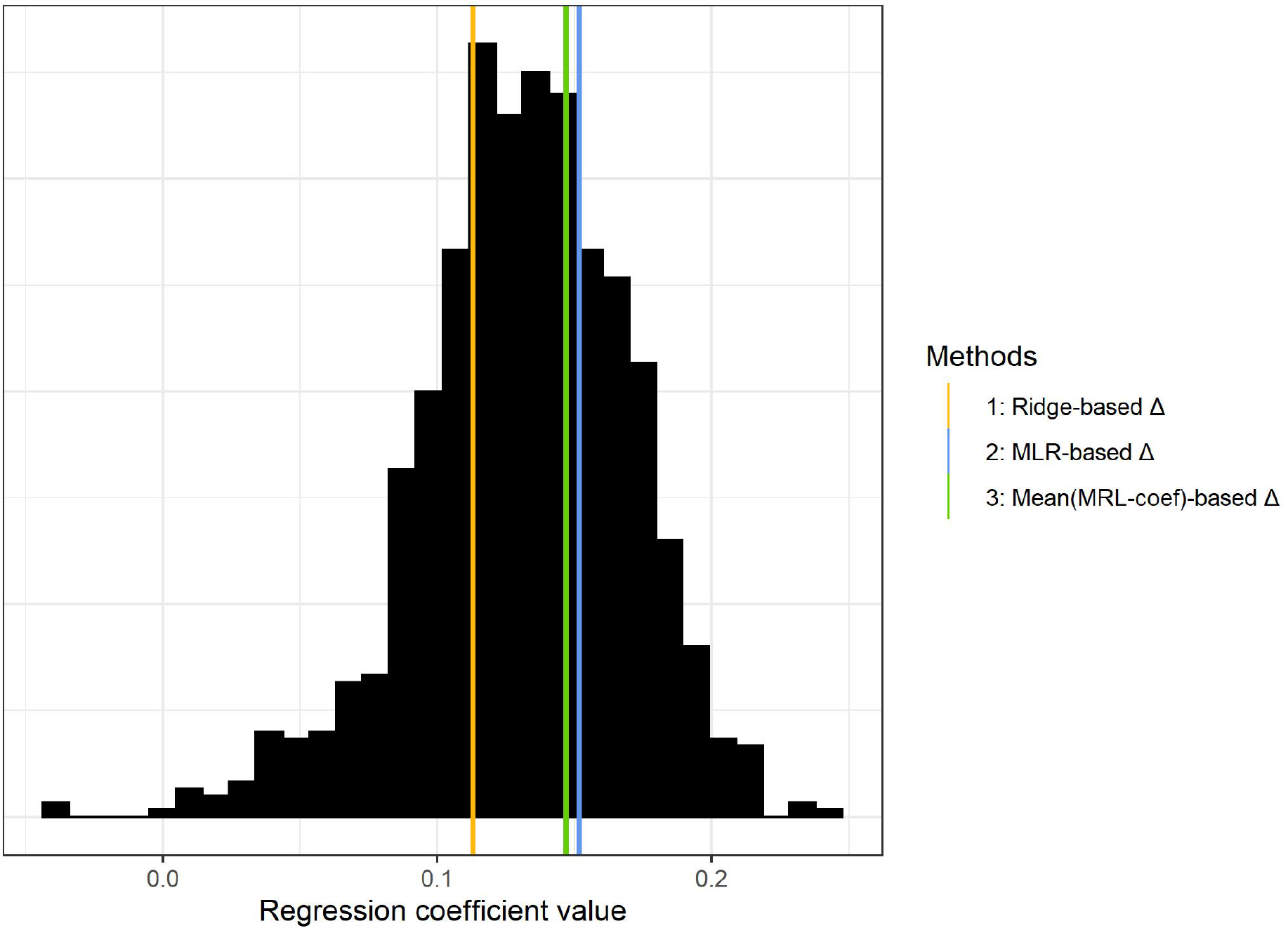
Regression coefficients (effect sizes) of the predicted Δ-values in a Cox PH model with time-to-mortality as the outcome, using the LLS data and 59 metabolic variables as predictor variables. The predicted Δ-values were calculated using 4 methods: using ridge regression (method 1), using multiple linear regression on a subset of metabolites (method 2), taking a linear combination where each metabolic variable was assigned the same weight (method 3), and taking a linear combination where each metabolic variable was assigned a weight randomly drawn from a standard uniform distribution, repeated 1,000 times (grey histogram, method 4).

**Figure 11:**
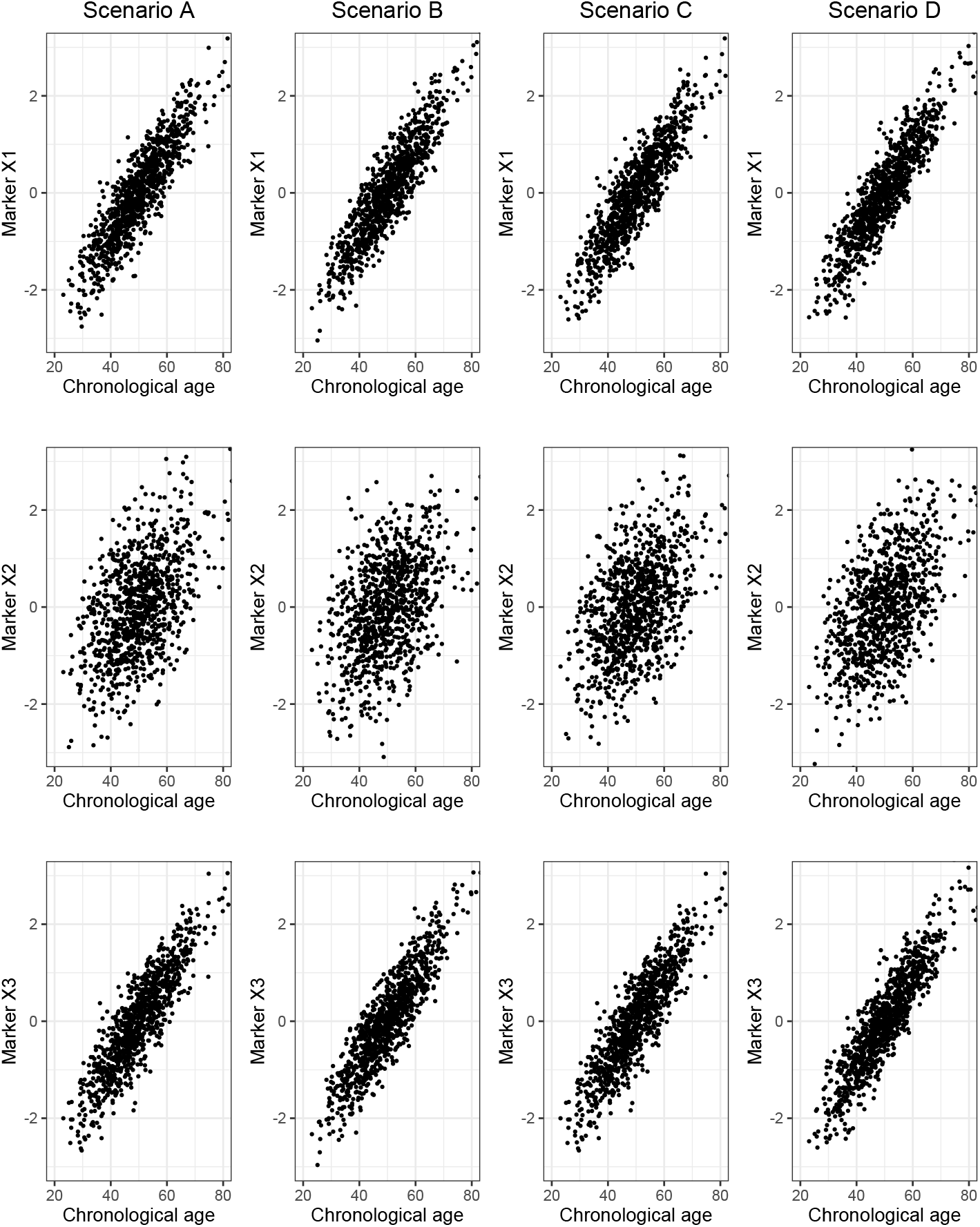
Scaled and centered markers *X*_1_, *X*_2_ and *X*_3_ plotted against chronological age for scenarios A (identical-association assumption holds), B (inverse association), C (no association) and D (identical-association assumption partially holds).

Although Figure 10 shows that predictions for rate of aging Δ obtained via ridge regression on 59 metabolic variables (method 1) are less strongly associated with mortality than predictions for Δ obtained using standard multiple linear regression on 26 metabolic variables (method 2), the *chronological* age predictions obtained with method 1 are more accurate than the ones obtained with method 2 (root-mean-square error method 1: 6.01, root-mean-square error method 2: 6.18). This exemplifies the biomarker paradox: improved chronological age predictions do not imply improved biological age predictions. In fact, after a certain point the association will weaken. We see that the better chronological age prediction performance of method 1 already results in a weaker association of Δ with prospective mortality.

Note that all coefficients in Figure 10 are positive. Since we kept the coefficient signs of method 2 for methods 3 and 4, it confirms our earlier assertion that if a marker is positively associated with chronological age, it is unlikely to be negatively associated with aging rate Δ (and vice versa). This explains why despite the suboptimality of cross-sectional methods, cross-sectional Δ-predictions have repeatedly been found to be associated with prospective mortality and other age-related outcomes [Marioni et al. 2015, Christiansen et al. 2016, Van Den Akker et al. 2020, Tanaka et al. 2020]—albeit (much) weaker than second-generation biological age predictors [Hillary et al. 2020, Maddock et al. 2020, McCrory et al. 2021, Kuiper et al. 2022]. The direction of the coefficients contains information regarding the signal. However, one must realize that unless the identical-association assumption (almost fully) holds, no more signal will be captured with cross-sectional methods than if markers would have been assigned weights at random.

## Discussion

We have shown that the most popular cross-sectional biological age predictors, where candidate markers of biological aging and chronological age are measured at a single point in time, all rely on the same underlying assumption: a candidate marker’s strength of association with chronological age should be directly indicative of its strength of association with the difference between biological and chronological age, also known as one’s aging rate Δ. We have called this assumption the identical-association assumption. We noted that there is no inherent statistical reason why a candidate marker’s association with chronological age *C* is indicative of its association with Δ: this depends on the biological context. Importantly, as we have proven, whether the identical-association assumption holds is untestable in a cross-sectional setting. As a consequence, one cannot distinguish true markers of biological age from false ones in such settings. A candidate marker can be correlated with chronological age but be completely uninformative of Δ. The opposite holds as well: a candidate marker may not be associated with chronological age, while being a true marker for Δ. We illustrated that unless chronological age and Δ are equally strongly associated with each marker, there is no guarantee that the size of the weights that a cross-sectional method assigns to candidate markers are informative of the underlying truth.

The identical-association assumption did not hold in the empirical data we considered. It should however be noted that we worked with a single real data set which is limited in size and scope. Our real data section is therefore primarily meant as an illustration of the potential practical consequences of constructing a cross-sectional biological age predictor if the identical-association assumption does not hold. It does not provide evidence for or against the extent to which this assumption holds in larger data sets or data sets with other types of candidate markers. Still, there is evidence that the identical-association assumption also does not hold in DNAm data: Levine et al. [2018] regressed a phenotypic age measure that captured differences in lifespan and healthspan on CpG-sites and found that the CpG-sites with the highest resulting weights did not correlate with chronological age at all.

Recently Nelson et al. [2020] addressed another important concern related to identification of aging markers based on cross-sectional data: mortality selection can bias the identification of markers, up to a point where cross-sectional analyses are less likely to identify true markers than if markers had been selected at random. While Nelson et al. [2020] state that this issue can be circumvented by only including markers that are known to be truly associated with mortality, in our second synthetic data example we illustrated that even in cases where all candidate markers are truly associated with biological age given chronological age, cross-sectional methods might not contribute either to selecting markers or to proving their validity.

We would like to stress that we do not claim that cross-sectional predictors of biological age cannot capture any signal. Although the identical-association assumption might not be realistic, for some (perhaps most) candidate markers the direction of a marker’s association with chronological age can still be informative. This also explains why many cross-sectional clocks were indeed found to be (weakly) correlated with various age-related outcomes [Marioni et al. 2015, Christiansen et al. 2016, Van Den Akker et al. 2020, Tanaka et al. 2020]: the sign of a candidate marker’s association with chronological age can be informative or uninformative of its association with rate of aging Δ, but it is unlikely to be be counter-informative. Hence, most cross-sectional methods can be expected to still capture some signal—but potentially not better than any other approach that in some naive or random way assigns weights to markers associated with chronological age. We do not reject the possibility that markers exist for which the identical-association assumption does hold. This assumption may or may not hold for different types of markers, but in a cross-sectional setting there is no way to tell.

Since there is no way in which the quality of a biological age predictor can be assessed using cross-sectional data alone, it follows that there is no way to optimize the quality of biological age predictions using cross-sectional data. Therefore, it is likely that biological age predictors based on cross-sectional data are highly suboptimal—they primarily capture signals related to chronological age, as also remarked by [Rutledge et al. 2022]—and that much better predictors could be constructed if researchers could work directly with longitudinal data.

This raises the question whether cross-sectional methods still have a place in the biological aging prediction landscape, or whether they should be abandoned completely in favor of methods that use longitudinal (time-to-mortality) data [Levine et al. 2018, Lu et al. 2019, Deelen et al. 2019]. By making the reasonable assumption that a higher biological age corresponds to a higher mortality risk, these time-to-mortality-based methods overcome the testability issue inherent to cross-sectional methods. The track record of these prospective mortality-trained methods in predicting various aging-related outcomes is indeed better than that of cross-sectional ones [Hillary et al. 2020, Maddock et al. 2020, McCrory et al. 2021, Kuiper et al. 2022]. Nevertheless, due to the relative abundance of cross-sectional data over longitudinal (time-to-event) data, cross-sectional predictors of biological age remain popular [Rutledge et al. 2022]. We think cross-sectional data can still play a role if the number of candidate markers is too high for to the limited sample size of the longitudinal data that is available and/or if there is little prior knowledge on the association between the candidate markers under consideration and aging rate Δ—which in this new era of high-dimensional omics-based aging clocks is quite a likely scenario. In such a case, cross-sectional data could be used to make a pre-selection of markers most strongly correlated with chronological age, as one might reasonably expect that at least part of these candidate markers will also be strongly correlated with Δ. Such a pre-selection does not have to be conducted in a multivariate way, but can be done per marker, as we did in our real data illustration.

Our view is that if longitudinal (aging-related outcome) data is available, methods using this information are to be preferred above cross-sectional ones to develop a biological age predictor. Depending on the extent to which the identical-association assumption holds in the data set under consideration, longitudinal methods might be preferred even if the sample size of the available longitudinal data is much smaller. Furthermore, we believe that the sizes of the coefficients of candidate markers obtained with cross-sectional methods should neither be used nor interpreted. If researchers do decide to develop an biological age predictor based on cross-sectional data only, they should be explicit about the underlying assumptions of the method they used and to what extent these assumptions are expected to hold.

## Data availability

All R-code used for the analyses in this paper is available in a public GitHub repository (https://github.com/marije-sluiskes/cross-sectional-bioage). Access to the individual-level data from the Leiden Longevity Study is restricted based on privacy regulations and informed consent of the participants. These data hence cannot be made publicly available. Data of the Leiden Longevity Study may be made available to researchers upon reasonable request to Eline Slagboom or Marian Beekman.

## Author contributions

MRG, HP, JJG and MHS developed the concept of this study. MB and PES collected the data used in this study. MHS and MRG performed the analyses. The first draft of the manuscript was written by MHS and revised by MRG, HP, JJG, PES and MB. All authors contributed significantly to this manuscript and approved the final version to be published.

## Acknowledgements

We thank Niels van den Berg, Erik van den Akker, Erik van Zwet and Bas Heijmans for the insightful discussions and helpful comments.

## Supplementary Materials

## Appendix A

This appendix contains the theoretical foundation underpinning our statement that in a cross-sectional setting the identical-association assumption is untestable.

Denote by *C* chronological age, by *B* biological age and by *X* a true marker of biological age given chronological age (*B|C*).

### Theorem.

*For every triplet* (*X, C, B*) *of continuous random variables, there exists another continuous random variable X*′ *such that* 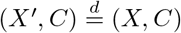 *and X*′ *is independent of B given C*.

*Proof*. Denote by *f* (*x, c, b*) the joint density of (*X, C, B*). Let

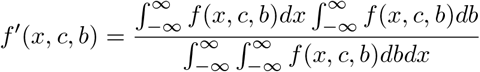

be the joint density of *X*′, *C, B*. As the integral of *f*’(*x, c, b*) over the entire space equals 1, this also constitutes a proper joint density. Moreover, this density is consistent with *f*, since

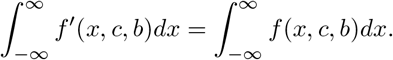

We have that 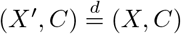, since

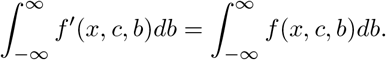

Further, we have that *X*′ is independent of *B* given *C* since 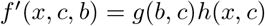 where

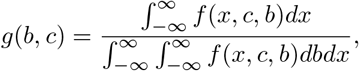

and 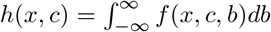.

Here we have considered a scenario with only one marker (*X*). In practice, researchers have many candidate markers to choose from, which are typically combined to a single biological age-metric. In that case, *X* or *X*′ can be viewed as the resulting biological age metrics. The theorem then asserts that we cannot distinguish between a good metric *X* and a bad metric *X*′.

We emphasize that the result of this theorem is independent of the method used to infer on biological age: such inference is impossible with any cross-sectional method.

## Appendix B

